# SPOILS OF WAR AND PEACE: ENEMY ADOPTION AND QUEEN-RIGHT COLONY FUSION FOLLOW COSTLY INTRASPECIFIC CONFLICT IN ACACIA ANTS

**DOI:** 10.1101/008904

**Keywords:** Crematogaster, Acacia drepanolobium, conflict costs, territorial aggression, worker relatedness, intraspecific slavery, colony fusion, Acacia ant, Pyrrhic victory

## Abstract

Intraspecific conflict over vital limited resources can lead to costly fights. How winners compensate for costs and minimize the threat of Pyrrhic victory is not well known. This study tracked the outcomes of experimentally induced field conflicts between highly territorial Acacia ant *Crematogaster mimosae* colonies using molecular genetics, and discovered that fatal fights significantly decrease within-colony worker relatedness. We find that reduced relatedness can be explained by colonies increasing worker number via 1) non-kin enemy adoption or 2) queen-right colony fusion. We hypothesize that incorporating non-kin enemies can speed recovery from conflict when resource defense is paramount. In the case of queen-right colony fusion, territorial defense benefits could outweigh fitness costs. We provide evidence that winners of colony fights have reduced worker forces to defend larger territories. Field assays indicate that post-fight colonies are more vulnerable to heavy browsing of host trees by mega-herbivores and takeover by competitors following conflict. We discuss the implications of our findings for ant colony cohesion and recognition systems.

---

Intraspecific conflict over resources can be costly. The evolution of assessment systems enables many adult competitors to settle disputes with minimal physical or energetic escalation [1,2] Still, intense combat does occur over particularly limited and valuable resources (e.g. mates and territory, but rarely food) [3–5]. Victory in such high-stakes battles can be decided through elaborate displays, or by violent contests [2] that continue until one opponent concedes from exhaustion or injury, or dies (wars of attrition [6]; ‘desperado effect’ [7]).

Winners benefit from defeating opponents by gaining access to contested resources. In addition, empirical studies widely document a positive feedback between successful fighting experiences and probability of victory in future contests, termed ‘the winner effect’ [8]. Yet by outlasting or dispatching conspecific contestants, winners are also theorized to accrue costs associated with escalation, including opportunity costs and loss of resource holding capacity [9,10]. As a result, winners may experience a diminished ability to defend themselves from predators and parasites, or to protect their gains in subsequent contests [11,12]. This possibility, that winners suffer increased vulnerability after engaging in fights, should be an important factor affecting potential costs and benefits of engaging or continuing in conspecific fights [13]; yet experimental tests are rare [5,14]. Furthermore, whether and how winners respond behaviorally to cope with fight costs and facilitate recovery remains largely unexplored. Here we investigate whether winners experience a window of vulnerability following costly fights, and if so how they might compensate to speed recovery from conflict.

Ants are compelling model organisms for this investigation for two main reasons. First, violent fights over territory are common [15]. Intruders are grasped or stung by defending workers, which often leads to dismemberment or death for both combatants. Second, the impact of colony conflicts to winners can be quantified in discrete units (i.e. individual workers killed). Most ant colonies exhibit reproductive division of labor where workers forgo direct fitness to rear immature workers and reproductives produced by the fertile queen(s). As a consequence of this ‘super-organism’ arrangement, loss of the queen(s) ultimately results in colony death [16]. Death of individual workers, however, need not be fatal to colonies. Instead, these losses can be viewed as the costs of conflict. Because colony size underlies victory in many ant species, significant reduction in worker number likely affects a winner colony’s ability to protect nest space and territory after fights. Hence, quantification of worker deaths is directly relevant to colony condition.

Do winners have tactics to recover from deficits incurred as a result of costly fighting? Models predict that protracted and dangerous fighting most often occurs when reproductive success or survival is at stake [4]. The mates or security (e.g. burrows, nest space etc.) that are the payoff of fight success cannot themselves be used to rebuild the winner’s physical condition post-conflict. One solution to rapidly replenish spent reserves can be cannibalization of defeated contestants. Some spiders, squirrels, moths, owls, and ants consume conspecifics after mortal combat for territory or shelter [17–19]. An alternative path to recovery uniquely available to social insects is rebuilding colony size by adopting a loser colony’s surviving brood and/or workers [20]. In doing so, energy lost in tissue conversion via consumption could be avoided, and time until viable workers are present is reduced (possibly even approaching zero). The challenge to this tactic is overcoming nest-mate recognition systems. Although often viewed as fortresses of cooperative relatives, ant colonies can be permeable to non-kin. Social and nest parasites invade host nests [21], unrelated queens found nests cooperatively [22], heterospecific colonies share nests [23], queenless colonies fuse [24] and some species, including invasive fire ants, reciprocally raid neighboring colonies for brood, creating genetically blended colonies [25]. Mechanisms enabling tolerance for non-kin within nests include host chemistry mimicry by usurpers, weak discrimination, recognition errors, signal mixing and environmental modification of recognition cues [24,26]. In the latter case, even contact with nest material can alter aggression patterns in ants [27]. Taken together there appears to be a strong possibility that individuals from defeated colonies can themselves become ‘spoils of war’, incorporated into post-conflict winner colonies regardless of relatedness to the usurping colony.

Through a series of field manipulations with the African Acacia-ant *Crematogaster mimosae* colonies we tested the hypotheses that intraspecific conflict for nest space on *Acacia drepanolobium* trees results in significant casualties for victor colonies and following these colony depletions, winners are less able to defend host trees against herbivores and competitors. We also use molecular markers to assess the outcome of conspecific fights and examine the impact of fights on the genetic composition of colonies. We predicted that territorial battles between conspecifics result in complete colony takeovers, and that post-conflict winner workforces are built back in part through the adoption of their non-kin enemies.

## METHODS

This study was conducted from July 2011 to March 2012 at the Mpala Research Centre in Laikipia, Kenya (37°53’ E, 0°17’ N). There as in many East African savannas, colonies of Acacia-ant *Crematogaster mimosae* battle elephants, giraffe, baboons, insect herbivores, conspecific and heterospecific Acacia-ant competitors (*C. nigriceps, C. sjostedi* and *Tetraponera penzigi*) to protect or maintain sole control of *Acacia drepanolobium* trees with which they are obligate mutualists [28]. Colonies gain additional domatia (swollen thorn nesting spaces) by initiating aggressive inter- and intra-specific wars to displace neighboring colonies from their host plants [28]. Territorial battles also occur when *A. drepanolobium* canopies grow together and when elephants topple trees into one another [29]. In high-density, monospecific stands of *A. drepanolobium* where >99% of trees are occupied by ants, up to 7.5% of resident colonies may lose host trees to other ant species over a 6-month period [30]. *C. mimosae* colonies are numerically dominant in the system (inhabiting 52% of trees [31]), suggesting substantial intraspecific conflict and turnover within this species, though exact rates are unknown.

Defeated colonies cede not only valuable domatia, but also their surviving immature and possibly even mature workers [20]. In this widely studied mutualism, there is no evidence of mixed species Crematogaster colonies forming after interspecific fighting. Striking morphological differences would make the presence of such colonies easy to observe. By contrast, in conspecific conflicts, the identity of winners and losers cannot be visually detected. To determine the consequences of conspecific conflict in this species we induced battles between neighboring C. mimosae colonies and used genes and behavior to track the consequences.

### Experimental Colony Selection

Mature *Crematogaster mimosae* colonies can be small and restricted to single host trees or form large multi-tree clusters with several queens and hundreds of thousands of individuals [30,32]. To ensure that colonies would fight in experimental battles and not retreat to auxiliary host trees, we chose only colonies inhabiting single trees with basal diameter of 32-68 mm (*X* + SE = 45.4±1.31). Colonies on trees of this size ranged from an estimated 2,697 to 9,870 workers (*X* + SE = 4999±255, calculated as domatia number * mean number of workers per gall, 68.5) [28].

Focal colony trees were all located close enough together that the canopies could be physically conjoined.

Single tree colonies were identified using reciprocal transplants of individual workers and watching for aggressive interactions with resident ants [28,33]. Latex gloves washed with 95% ethanol prevented between-trial chemical contamination of individuals. To aid observation of fast moving transplanted workers, we applied florescent powder (Day-Glo Color Corp., Cleveland, OH) to the thorax of trial ants. We treated individuals of the resident colony in the same manner to act as procedural controls. If non-resident ants were quickly attacked (but resident controls were not) we inferred that trees belonged to separate colonies [28]. We repeated this method on neighboring trees up to 8m away to confirm experimental colony monodomy. Each colony within the pair was identified with metal tags as either A or B with a shared Fight ID number (Table 1).

**Table 1.** Sampling design, colony relationships, and experimental fight outcomes for *Crematogaster mimosae*

| Fight pair ID | sample n (A, B, 2mo, 8mo)<sup>a</sup> | # loci (A, B, 2mo, 8mo)<sup>b</sup> | # queens (A, B, 2mo, 8mo) | Winner (behav assay) | Winner (molecular data) | Adopt enemies<sup>c</sup> | Prop adoptees (at 2mo) | Between colony relatedness (A vs B)<sup>d</sup> | A and B genotypes present after 8mo? <sup>e</sup> |
| --- | --- | --- | --- | --- | --- | --- | --- | --- | --- |
| 4 | 8, 8, 48, 22 | 17, 17, 4, 17 | 1,1,1,1 | unclear | A | no | 0 | 0 | no |
| 7 | 8, 8, 44, 24 | 17, 17, 5, 17 | 1,1,1,1 | unclear | B | no | 0 | 0 | no |
| 1 | 8, 8, 47, 24 | 17, 17, 4, 17 | 1,1,2,1 | unclear | B | yes | 0.21 | 0 | no |
| 2 | 8, 8, 46, 23 | 17, 17, 4, 17 | 1,2,3,1 | A | A | yes ★ | 0.17 | 0.08 | no |
| 5 | 8, 8, 47, 23 | 17, 17, 4, 17 | 1,1,2,1 | A | A | yes | 0.44 | 0.04 | no |
| 6 | 8, 8, 47, 23 | 17, 17, 3, 17 | 1,1,2,1 | A | A | yes | 0.04 | 0 | no |
| 8 | 8, 8, 45, 24 | 17, 17, 4, 17 | 1,1,2,1 | B | B | yes | 0.04 | 0 | no |
| 9 | 8, 8, 47, 23 | 17, 17, 5, 17 | 1,2,3,2 | unclear | ? | ° | fusion | 0.04 | yes |
| 3 | 8, 8, 46, 21 | 17, 17, 4, 17 | 1,1,2,3 | A | ? | unclear # | fusion | 0.03 | yes |

A = individuals collected from A tree (Pre-fight)
B = individuals collected from B tree (Pre-fight)
2mo = individuals from joined A+B trees (2 months after fight)
8mo = individuals from joined A+B trees (8 months after fight)
<sup>a</sup> n - number of individuals (indv) genotyped from each period (A, B, 2mo all indv = workers; 8mo indv = pupae or larvae)
<sup>b</sup> 2mo loci restricted to those with diagnostic alleles differentiating A and B genotypes
<sup>c</sup> Proportion of sample individuals assigned as full sibs with loser colony genotype with (inclusion/exclusion probability > 0.99).
Exceptions-
★ ID 2 - half sibs with B = 7 indv (p > 0.95)

#### Manipulations

Between 14-July and 04-August 2011, we induced fights between colonies by tying the canopies of the two experimental trees together with wire. *Acacia drepanolobium* stems are flexible and tolerated bending. Canopies remained connected for 8 months after fights.

Immediately prior to fights we collected three healthy domatia filled with live workers and brood from each tree. These ants were kept contained and isolated in the lab, and fed on a diet of sugar water and tuna. To determine the identity of the winning colony, live individuals (*N* = 2-4 from each of the pre-fight A and B colonies) were returned to the field and placed on the main stems of both trees on day 6 after fights and observed in the manner of reciprocal transplants described above. Winner and loser colonies were determined by the combined outcome of these behavioral experiments and molecular genetics (below).

#### Costs of Fighting to Winners

During territorial battles, workers from each colony engage in fights to the death, with larger colonies the more likely victor [28]. Colony fights are expected to produce heavy winner casualties. To quantify these costs we placed large plastic tarps secured at ground level between some paired fight trees (*N* = 7) and collected workers that fell from host plants when canopies were experimentally joined and fighting commenced. Dead and injured workers on tarps were collected every 24 hours (if fights lasted more than one day). Non-ant debris was removed and remaining workers were dried and weighed (Mettler Electronic Balance). The weight of 10 intact workers/10 gave an average individual worker mass for each colony. Total casualties were estimated from the mass of tarp casualties. Our casualty values are likely underestimated as wind removed some dead ants from tarps in the field.

To parse winner and loser colony contribution to total casualties, we genotyped (N=15-16) individuals collected from tarps of 5 of these 7 fight pairs and matched them to their respective colonies following molecular protocols (below). We calculated the cost to winner colonies as the total worker loss and proportion of the initial colony lost.

#### Vulnerability Associated with Fighting

Newly acquired territory (as well as original host trees) may be precariously defended by a diminished worker force after fights, and at risk from attack by other space-limited neighbors. To assess vulnerability, we selected single tree colonies similar in size to experimental fight trees as controls (N = 10 each for controls and experiment fights; Welch’s t test for tree diameter difference between groups *t*_13, 0_ = −1.74, *P* = 0.105).

We examined changes in colony response to simulated large mammalian herbivore browsing using methods modified from [34]. Two observers carefully approached trees. Each visually identified an isolated branch with new growth, and one swollen thorn domatia within 15 cm of the tip. Disposable ‘mitts’ crafted prior to fieldwork (two sheets of paper towel 11”x6” folded over by 1 inch and taped along 3 sides) were placed on each surveyor’s right hand. With a leather-gloved left hand, focal branches were simultaneously raked 3 times and then enveloped by the mitt. Worker ants swarming on mitts after 30 seconds were collected along with the mitt into a sealed bag. Bags were frozen and worker contents subsequently counted. Surveys occurred on each of the 3 days before experimental fights were induced (within tree replication *N* = 6; 2 surveyors x 3 days), and then again beginning 6 days after paired fights concluded.

Tree main stems are a primary access point for host invasion by ant competitors [33,35], We used photographs taken 3 days before fights and again 6 days after to assess changes in colony defense of stems on the simulated herbivory trees from above (n=5 photos from each period, 2 morning and 3 afternoon). Macro-digital photos always captured the south-facing plane of stems and adjacent size standard ruler. We recorded the number of ant heads visible in the frame from ground up to 10 cm.

Change in average response within colonies between controls and experimental fights were analyzed using *t*-tests in JMP 8.0.

#### Colony Relatedness and Enemy Adoption Sampling / collection methods

Individuals from *N* = 18 experimental colonies were analyzed using molecular techniques. Before trees were tied together we collected pre-fight workers from each experimental colony into a collecting vial (70% ethanol) in the field.

For post-fight collections, we collected 5 domatia from each experimental fight combined tree system on Sept 15, 2011 (hereafter 2mo) and March 22, 2012 (hereafter 8mo). Domatia were frozen, then opened and the contents (mature and immature ants) were pooled into vials with 70% ethanol. For pre-fight samples as well as 2mo samples, only intact adult workers were genotyped. Individuals were inspected under a dissecting microscope to ensure they had no appendages missing (indicating they were alive at the time of collection and not cached casualties or emerged workers that were recently killed) and to exclude body parts of other individuals. For 8mo samples we genotyped only immature ants (nearly all worker pupae, but for colonies with < 24 pupae we extracted DNA from large larvae or male pupae). Immature ants collected 8 months after fights are unlikely to have been present at the time of conflict (Development time - Appendix 1). Outcomes where all 8mo samples belong exclusively to one colony suggest that the takeover was complete and resulted in the loser queen’s death. Alternatively, if genotypes matching both Pre-fight colonies could be found among the immatures, an incomplete takeover is indicated (no loser can be identified because both queens survived and continued to contribute to worker production). Finally, novel genotypes that could not be explained by different fathers but the same mother would suggest that a new queen was present within the colony. This could occur though secondary takeover by non-relatives, or possibly via the emergence of reproductive daughter queens [32]. For colonies in each fight pair, sample sizes are listed in Table 1 but roughly follow: 8 workers from each colony prior to fights (Pre-fight), then 48 workers and 24 immature ants after 2 months and 8 months respectively.

#### Lab protocols

Individuals were extracted either using Qiagen DNA easy kits or Qiagen Puregene extraction techniques. We genotyped Pre-fight and 8mo individuals using 17 microsatellite loci (PCR protocols described in [36]).

For 2mo samples, where 48 individuals from each fight pair were analyzed, we first determined which loci contained alleles that could distinguish between fight pair colonies (from Pre-fight analysis results). We selected the 3-5 loci that differed most in frequency between colonies, and included only primers from those loci in the PCR reaction. If individuals could not be definitively assigned with the subset of primers, they were re-run with more loci. PCR products were run on a Capillary Electrophoresis Genetic Analyzer (an ABI Prism 3130) at UCDNA Sequencing Facility at UC Davis and analyzed using GeneScan software (Applied Biosystems, Carlsbad, CA, USA). Fragment data were visualized and scored using STRand Version 2.3.69 [37].

#### Analysis of genetic data

Parentage for each worker was reconstructed using a maximum likelihood approach implemented in COLONY v2.0.1.8 [38]. Null alleles and scoring errors were accounted for using a 0.05 default error rate at all loci, and no *a priori* relationships were assumed. Data for all individuals (from all fight pairs at all times, *N* = 765) were combined for analysis. Individuals were separated into full sibling or half sibling families, with associated probabilities of inclusion and exclusion for each individual (Table 1). Queen number was also estimated, and the identity of the maternal and paternal lineages at each sampling period were reconstructed.

Allele frequencies obtained from COLONY analysis (above) were used to calculate relatedness in COANCESTRY v 1.0.0.1 [39]. Relatedness values between Pre-fight colonies were based on all genetic data from 17 loci. Because 2mo individuals were genotyped at a reduced number of loci, relatedness estimates within individual winning colonies across all time points (Figure 2) were based only on data from those restricted 3-5 diagnostic loci. For fight pairs where genetic reconstruction did not reveal a distinct winner (ID3 and ID9), the Pre-fight relatedness (Figure 2) used for statistical analysis is the mean of the among-individual, within-colony relatedness for both Colony A and B. For each fight pair, individuals were grouped by their sampling origin, Colony A Pre-fight, Colony B Pre-fight, 2mo, 8mo, and compared to all individuals within the group to produce average within-colony relatedness at each time period. We report Triadic Maximum Likelihood estimators of relatedness coefficients, (TrioML) because values are restricted to fall from 0-1, making interpretation intuitive. In contrast to other pairwise relatedness estimators, this measure uses a third reference individual to help minimize error [40]. To test for effects of fighting on relatedness within colonies through time, we compared the TrioML relatedness values across Pre-fight, 2mo, and 8mo sampling periods using standard least squares regression, accounting for repeated measures by including Fight ID as a random effect. Since TrioML relatedness coefficients are bounded at 0 and 1, we performed arcsine square root transformations prior to analysis to better meet the assumption of normality

**Figure 2.**
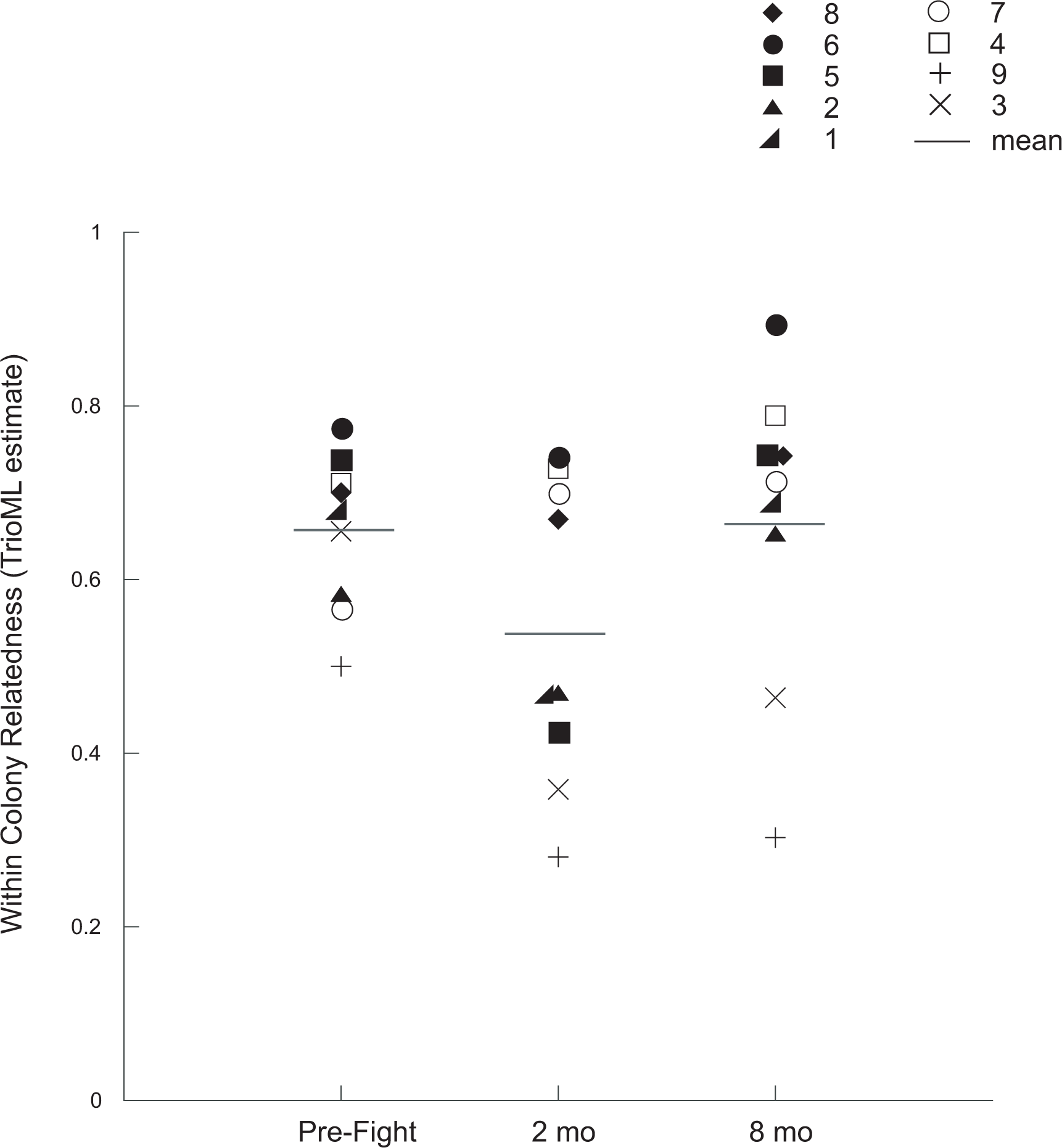
Average within colony relatedness over time for each fight manipulation. Pre-fight relatedness values are from fight winner colonies (except for ID3 and ID9). For these two colonies where no definitive loser could be identified, values are an average of within Pre-fight Colony A and B relatedness. Workers collected 2 months after fights were significantly less related on average to their nestmate contemporaries than workers sampled before fights and brood sampled 8 months after (*P* <0.0001). Winner colonies ID4 and ID7 (open symbols) contained no loser genotypes 2mo after fights. Solid symbols identify colonies with enemy adoptees (loser genotypes present at 2mo but not at 8mo). Line symbols identify colonies with no definitive fight winner where colony fusion is inferred (both pre-fight colony genotypes are present in workers at 2mo and brood at 8mo).

**Figure 3.**
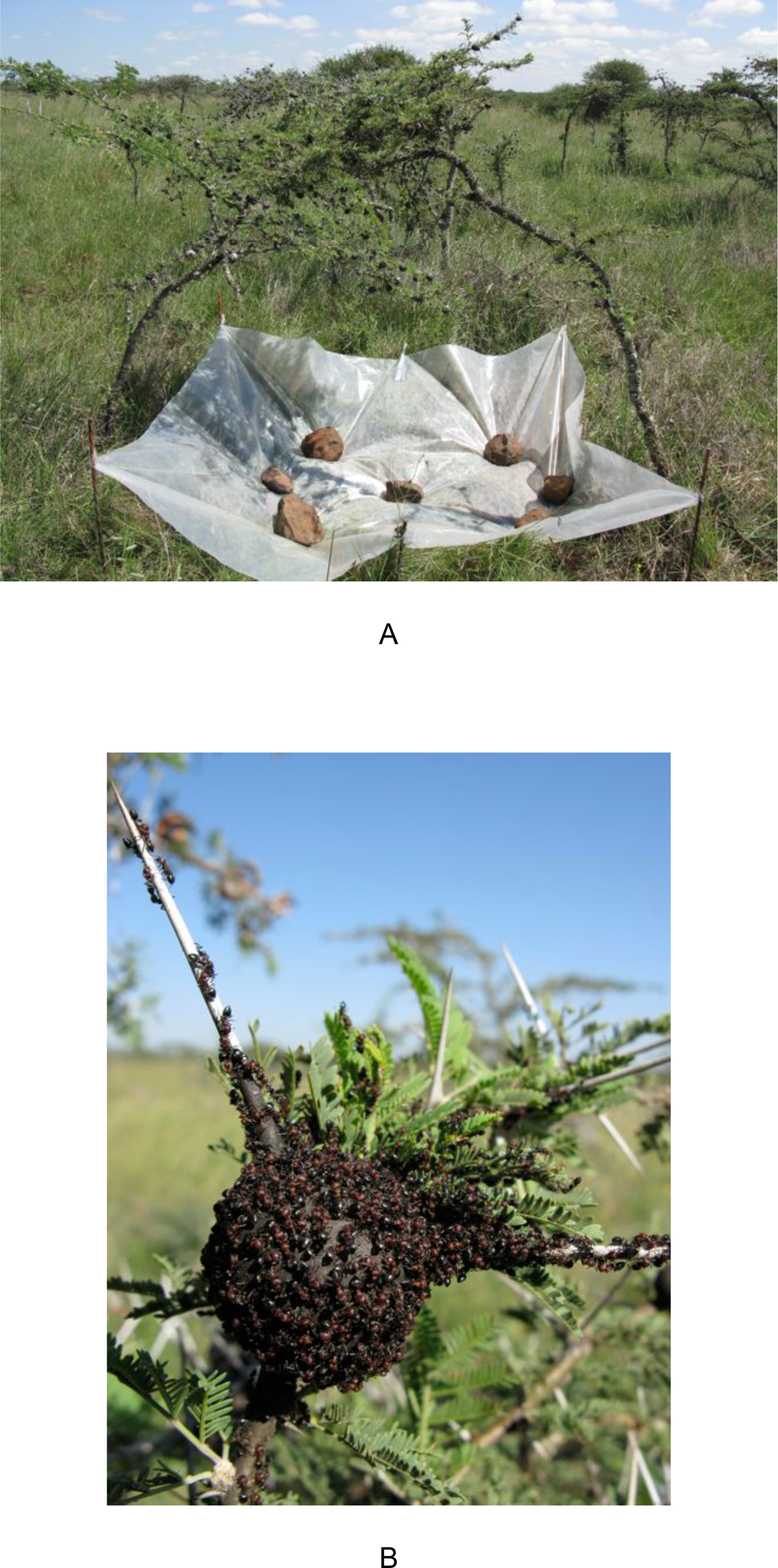
Images of experimental fights. Conflict induced by joining host tree canopies of separate colonies (A) with a tarp to catch casualties. (B) *Crematogaster mimosae* battle in progress on the surface of a contested domatia.

## RESULTS

### Costs of Fighting for Winners

Estimated worker losses in fights between colonies ranged from 390-10 073 individuals (*N* = 7 colonies, *X* +SE = 5,405+1,313 individuals). A subset of these casualties were genotyped and matched as full or half siblings to Colony A or B pre-fight (Pre-fight) samples, all with a probability of assignment >0.94 in maximum likelihood sibship configurations from COLONY. Winning colonies experienced high costs to fighting, with 19-56% (*X* +SE = 39 +7%, *N* = 5) of the casualties collected for each fight pair belonging to the victors. Successful colonies therefore lost on average 1/3 (*X* +SE = 34+11% *N* = 5) of their initial worker force.

#### Vulnerability

Following fights, winner colony territories nearly doubled (proportion of domatia pre- versus post-fight; *X* +SE = 1.9 + 0.08 *N* = 14) and defense of host trees declined significantly. Canopy defense dropped by more than half compared to Pre-fight levels as fewer ants from winner colonies responded to simulated branch herbivory than from control colonies (*t* test: *t*_18_ = −3.12, *P* = 0.006, Fig 1-A). Winner protection of stem access points also fell, with a marginally significant difference between treatment and controls (*t* test : *t*_18_ = −1.95, *P* = 0.067 Figure 1-B).

**Figure 1.**
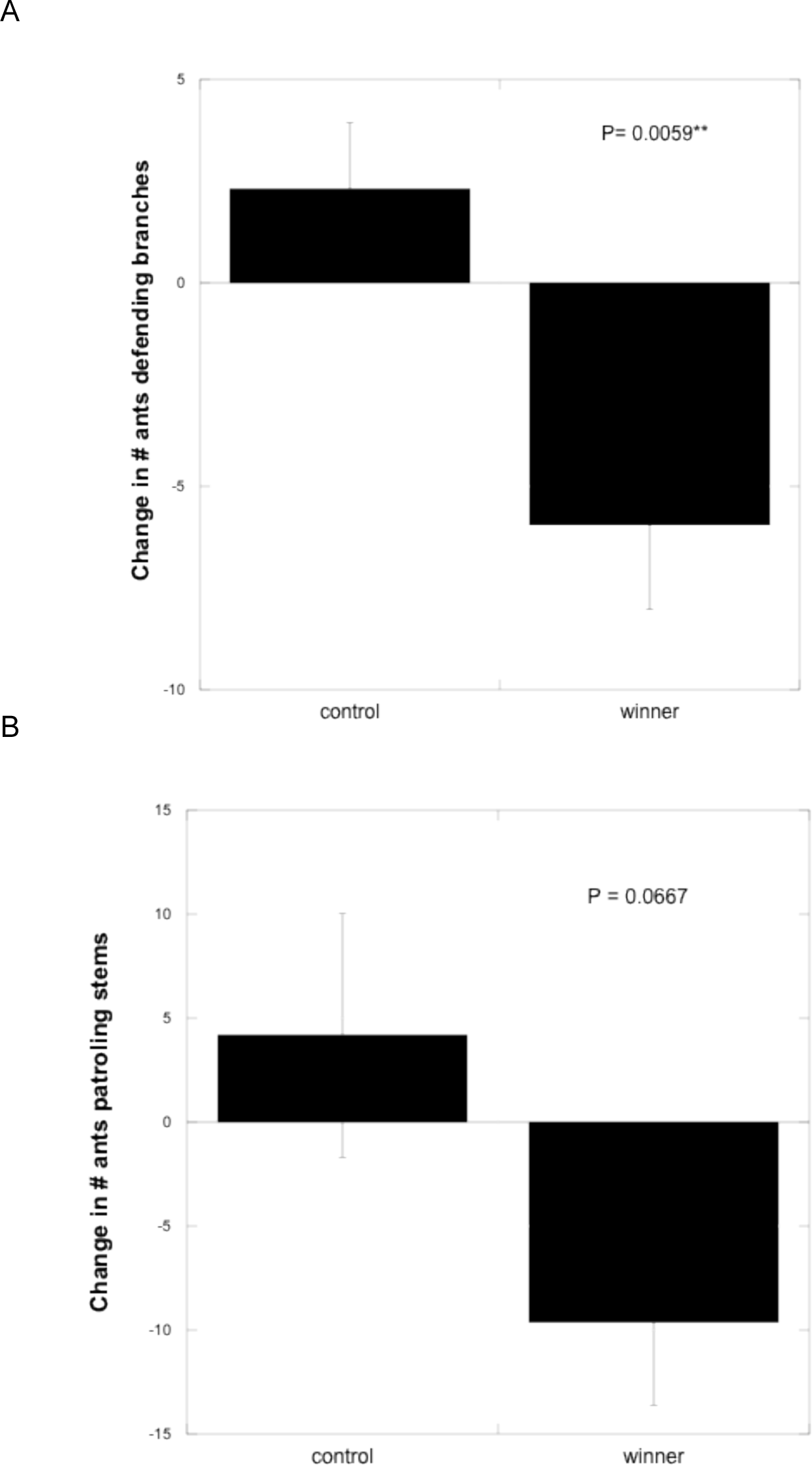
Vulnerability following conspecific conflict. Winner colonies decreased defense after battles. Mean +SE change in the number of ants guarding host trees in canopies (A) and on main stems (B) relative to pre-fight levels. Unmanipulated controls did not show a decrease in defense over the same period.

### Colony Genetic Structure

Pre-fight relatedness between colonies was low for all 9 fight pairs analyzed using molecular markers (TrioML estimate of *r* < 0.08 for all pairs, Table 1). Sixteen of the 18 colonies were determined to contain full and half sib workers produced by a single queen and two Pre-fight colonies were determined to include workers produced by multiple queens (Table 1).

We infer three different outcomes from molecular analysis of fights; complete rejection of non-kin, enemy adoption, and fusion. For fight pair ID’s 7 and 4, all post-fight individuals (*N* = 68–70 from 2mo and 8mo) were assigned as full siblings with individuals from only one Pre-fight colony (complete rejection of non-kin). The remaining 7 pairs contained workers at 2 months after fights that were matched as full siblings with individuals from both Pre-fight colonies Consistent with enemy adoption, at 8mo, genotyped brood (*N* = 23 or 24) from 5 of these 7 pairs were assigned to only one of the Pre-fight colonies. For these colonies, molecular data suggests a single queen-right colony succeeded in conflict and subsequently included unrelated workers, but confirmed that loser queen(s) were either no longer present or no longer contributing brood to the colony at 8 months. Surprisingly, for fight ID’s 3 and 9, genotyped brood at 8mo were full siblings with individuals from both Pre-fight colonies, indicating that one colony did not completely overtake the other and that both Pre-fight queens were present and producing offspring. Furthermore, 16 individuals (76%) of 8mo samples from fight ID 3 were classified by COLONY as full sibs with each other (probability inclusion /exclusion >0.99) but were inferred to be the offspring of a novel maternal genotype. Overall within-colony worker relatedness decreased significantly from Pre-fight (*X* +SE = 0.72 +0.03) to 2mo samples (0.52 +0.08;

Linear mixed model: *R*^2^ =.80, *P* <.0001; TukeyHSD for Pre-fight, 2mo comparison: *P* = 0.008). Average relatedness for brood found within each colony at 8 months after fights *X* +SE = 0.68 +0.08) was similar to Pre-fight worker relatedness (Tukey HSD for Pre-fight, 8mo comparison: *P* = 0.760) indicative of restoration of pre-fight relatedness conditions at a colony level (Figure 2).

## DISCUSSION

### Worker Losses and Vulnerability

Are territorial fights between *C. mimosae* colonies costly to winners, and does conflict weaken the victor’s ability to defend resources? We found that battle casualties reduce winner colony size by 1/3 on average, and as much as 2/3. In contrast to investigations highlighting advantages to combat victory beyond resource acquisition (e.g. probability of success in future conflicts [33,35] and better health [41]), we find that after successful fights, winning ant colonies suffer from a window of vulnerability. *C. mimosae* winners are spread over twice the territory (initial plus newly gained domatia) and appear susceptible to a loss of territory value (e.g. removal of extrafloral nectaries, domatia, and reduced tree growth) from mammal browsing and insect herbivory. Previous manipulations of ant abundances on *A. drepanolobium* trees reveal a negative relationship between colony size and branch damage by elephants and beetles [42,43]. The actively growing shoot tips favored by large herbivores are also the site of carbohydrate-rich extrafloral nectar production, which colonies rely on to fuel activity and feed developing larvae [31,44]. Less than one week after experimentally induced wars, colony defense of host tree canopies dropped by 66% compared to pre-fight levels (Figure 1). Simulated browsing of these resources incited no defense on 3.5x the number of branches sampled after fights as compared to before. Inability to protect tree-based energy sources should hinder a colony’s ability to produce and sustain workers. Additionally, activity on tree trunks fell by 62% after fights, and we found a complete absence of workers patrolling trunks in nearly 5x as many post-fight observation periods as pre-fight observation periods. It is possible that nest-limited neighbors discover the diminished resource holding capacity of winners either via territory scouts [45] or by eavesdropping on the pungent alarm pheromones released during combat [46,47]. Neighbors may then apply the information gained through monitoring to target weakened competitors [48]. In a system where colony size underlies competitive success [28], we document decreased worker number and defense of hosts by winners. We hypothesize that public battles fought to gain territory may subject victor colonies to increased risk of attacks and territory loss [12,49].

### Non-kin Adoption During Recovery

Are winner colonies built back though the adoption of losers? We found that following fights, former non-kin enemies coexist within shared nests. In 56% of the induced fights we analyzed genetically, despite prior lethal aggression between competitors, post-fight colonies contained live workers that were full siblings with the pre-fight loser colony (Table 1). In these cases, losers represented an estimated 4-44% of post-conflict colonies’ workforce. This integration of losers in these cases was not consistent with queen-right (both queens present) colony fusion because no brood developing within winner nests matched loser genotypes at 8 months after fights. We conclude that for these five colonies, loser queens were either killed or escaped during fights. We further infer that their offspring (undeveloped brood, and possibly surviving workers) are adopted by the victors, and act as an ephemeral resource for the winner colony. Genetic similarity between winner and loser colonies does not explain variation in the incorporation of non-colony members as all averaged pair-wise relatedness values between fighting colonies were very low (relatedness coefficient *x* < 0.08 for all colony pair comparisons, *X* +SE = 0.02 +0.06; 1^st^ cousins should exhibit *r values* of ∼0.25; Table 1).

We estimate that 1,700-5,550 loser brood could remain within domatia after experimental takeovers. This calculation is based on the average number of brood per domatia (*X* = 37.3 from [50]) and the number of domatia on experimental trees (*N* = 48-150). Since colonies in fighting pairs were estimated to each contain a similar number of brood prior to conflict, adoption of all loser brood would provide an instantaneous near doubling of new workers emerging within the winner colony. Randomly selected *C. mimosae* pupae reared in the lab become workers in an average of *X* +SE = 8 +0.74 days, and well-developed larvae pupate after an average of X +SE = 12 +0.72 days (Appendix 1). Pupae can metamorphose into fully formed workers with no tending or nutritional input from adults (Appendix 1). Although complete worker development time (egg to adult) is unavailable for this species, Argentine ants (*Linepithema humile*) display pupal development rates that are similar to those observed for *C. mimosae* and require an average of 63 days from oviposition to worker emergence [51]. Consequently, feeding loser brood to egg-laying queens or to larvae instead of adopting them directly would create a months-long payoff lag. Retaining rather than consuming loser brood may further boost colony size by stimulating the surviving queen’s egg production, as is seen in *Oecophylla* weaver ant colonies that are experimentally augmented with non-kin pupae [52]. Adoption of abundant non-kin brood is thus a more efficient way (in terms of both time and energy) of converting loser individuals into valuable workers than a potential alternative – cannibalism.

Sterile conspecific workers laboring for the fitness benefit of an unrelated queen and/or colony - historically termed intraspecific slaves (but see [53]) – have few documented examples in wild ant colonies [15,54]. This study represents a rare example of this phenomenon, and the first evidence that non-kin enemy adoption can be triggered experimentally in nature via conflicts between large territorial colonies. Previous descriptions of natural non-kin conspecific adoption/enslavement in ants come from species that are close relatives to facultative and obligate interspecific slave-making taxa [15,55,56]. *Crematogaster*, a species-rich genus (476 known species - [57]), has no known obligate slavemakers [15,21,58] and diverged from known obligate slave-making species > 80 mya [59]. Like many other obligate plant ants and cavity nesters, *Crematogaster* spp. are known to compete strongly with conspecifics for nest space [60] and invade the domatia of heterospecific neighbors [35]. Conspecific usurpation, though difficult to detect in nature, has been predicted [55] and observed ([28], K. Rudolph pers. obs.). Our findings suggest that non-kin adoption associated with territorial battles could be an overlooked phenomenon, and potentially widespread in ants that engage in conflicts over nest sites or foraging grounds.

### Additional Fight Outcomes

Experimentally induced fights had multiple distinct outcomes. While five winner colonies adopted non-kin orphans, two fights unexpectedly resulted in queen-right colony fusions (Table 1). In fight ID’s 3 and 9, genetic data indicates that both Colony A and B queens were alive and producing brood within a shared tree canopy 8 months after fights (Table 1). A unique feature of *C. mimosae* seems to be that intense intraspecific aggression with extensive mortality can rapidly give way to tolerance. In other documented cases of non-kin mergers, worker interactions are rarely characterized by high aggression or lethality [24,25,54,55,61–65]. Yet we find that in *C. mimosae* colonies mortal combat transitions to coexistence over the course of hours (active fighting never lasted for more than 48 hours, and most wars resolved in less than 12 hours). Not only does aggression toward brood and callow workers cease, but it seems worker-worker attacks do as well. It appears unlikely in the case of colony fusions that queens could survive fights without some or many of their adult defenders also persisting.

We do not yet know how de-escalation between fighting colonies may proceed. It is possible that for conflicts between social insects generally, and especially colonies with weak size asymmetry, assessment of fighting ability is difficult [66]. Fights may escalate because the superior competitor cannot be readily determined (failure of mutual assessment, as discussed in [49], or because contested resources (e.g. nest space) are essential for colony survival (i.e. there is no assessment; [67]. However, information indicating a growing cost of conflict (e.g. fight duration) may induce de-escalation behavior in workers, and protect against Pyrrhic victory. Reciprocal de-escalation in both colonies could result in such a truce. Non-kin colony cooperation in times of vulnerability has precedent in ants (e.g. joint colony founding ([22] and queenless colony fusion [24]) but has been previously unreported in large, mature colonies as a result of conflict. Regardless of the de-escalation mechanism, ultimate coexistence appears impossible without strong temporal plasticity in templates of recognition or chemical cues among adults.

Uncovering the patterns and mechanisms of recognition within and between social insect colonies has long been an area of interest for biologists. Two primary concerns are how and when the signals used in nestmate discrimination are acquired. Chemical signals and perception underlying worker exclusion or acceptance in a colony are not solely inherited; there can be ecological and environmental effects on both [26,27,68]. Cooperation among workers can be mediated by queen pheromones and/or learned based on differences in cuticular hydrocarbons (CHC). These chemical signatures can be modified by the environment and spread among individuals. In our cases of non-kin adoption, winners’ templates for tolerance appear to change as a result of fighting. Non-kin losers may retain cues that winners detect as distinct but overlook to enable acceptance of more individuals into the colony (active adoption/ tolerance; [65]. Alternatively, through the extensive physical contact involved in fighting, individuals may blend and dilute CHC signals, making non-kin indistinguishable from one another (passive affiliation; [20,24,69]. We suggest this system may be fertile ground for future studies examining how colony chemistry is altered by conflict, especially as host tree takeovers are frequent.

Our third outcome showed that two winner colonies did not adopt non-kin (at least in numbers appreciable in our samples, Table 1). During fights representing each of our three outcomes discussed above, we occasionally witnessed advancing workers ejecting brood from their opponent’s domatia. If all loser workers were eliminated in battle and this brood ejection behavior continued, it could explain the lack of non-kin in all post fight samples for ID’s 5 and 7. The findings of complete rejection, as well as temporary rejection followed by acceptance of non-kin in this study, underscore the tension in colonies between the benefits of incorporating non-kin versus the threat that non-kin could represent to colony cohesion. Further exploration could help determine whether reduced relatedness among nestmates following conflict has unexpected consequences for colony function and the extent of recognition plasticity. For example, do adoptees perform work within winner colonies and are some colonies’ signatures fundamentally incompatible with others?

### Conclusions

Our findings represent a rare experimental quantification of the costs and consequences of escalated fighting for winners in their natural environment. Application of molecular analysis to a behavioral study exposed the leakiness of colony boundaries in *C. mimosae.* Through field manipulations, we produced a pattern long inferred by other researchers [17,55,56,61,70] that ant conflicts over territory predictably decrease within-colony relatedness (via non-kin enemy adoption and colony fusion Fig 2), and that colony cohesion appears robust to this perturbation. Importantly, this non-kin affiliation occurs within large mature colonies, not recently founded ones [71] A study of conflict in wood ants showed that violent wars fought to expand territory produced casualties that were fed to developing larvae [72]. Our work points to a different, potentially underappreciated source of profit for colonies that succeed in conflict. After costly contests, *C. mimosae* winners at times gain not only valuable new host trees but also living spoils of war in the form of non-kin adoptees that provide victors with an accelerated means to colony size recovery.

## ACKNOWLEDGEMENTS

This study was carried out with permission from the Kenyan Government (Permit # NCST/5/002/R/483 to KPR). Funding from the University of Florida, Explorers Club, Sigma Xi, and the American Philosophical Society. Many thanks to intrepid field assistants Antony Eshwa, Alfred Inzauri, James Ekiru, and John Lemboi and to all the staff of the Mpala Research Centre. Sincerest thanks to Maureen Stanton, Rick Grosberg and Brenda Cameron for assistance with molecular work. Manuscript benefited tremendously from comments made by Lauryn Benedict, Neil Tstsui, Todd Palmer, Jake Goheen, Jane Brockman, Christine Miller, Dan Doak, Brian Silliman.

